# Phosphoarginine modulates oligomerization and repressor activity of mycobacterial ClpC2

**DOI:** 10.64898/2026.06.30.735635

**Authors:** Henry R. Anderson, Pratistha Kandel, Emmanuel C. Ogbonna, Karl R. Schmitz

## Abstract

Phosphoarginine (pArg) modifications direct proteins for proteolytic destruction by ClpC1P1P2, an essential mycobacterial protease that has emerged as a promising antibacterial drug target against *Mycobacterium tuberculosis*. The broader regulatory landscape surrounding pArg is poorly understood. Here, we establish a mechanistic connection between pArg binding and the activity of ClpC2, a non-proteolytic transcriptional repressor with homology to the ClpC1 N-terminal domain. Biophysical studies reveal that ClpC2 forms concentration-dependent higher-order oligomers that bind cooperatively to operator sequences in the *clpC2* promoter. A high-resolution crystal structure of the *Streptomyces thermoviolaceus* ClpC2 C-terminal domain reveals a conserved dimerization interface mediated by a C-terminal helix, which is sterically disrupted by pArg binding. Consequently, we find that binding of pArg, as well as some ClpC1-targeting antibiotics, disrupts ClpC2 oligomerization, dissociates ClpC2 from its operator DNA, and relieves transcriptional repression *in vitro*. Moreover, comparative analysis of *clpC2* promoters with single versus dual operator sites predicts differences in regulatory sensitivity across mycobacterial species. Together, these findings establish ClpC2 as a pArg-responsive sensor capable of mechanistically linking elevated pArg levels to downstream transcriptional regulation.

## INTRODUCTION

Tuberculosis has been a major driver of human mortality throughout the history of civilization. The causative agent, *Mycobacterium tuberculosis*, infects a quarter of the world’s population and causes >1 million deaths annually [1], more than any other single pathogen. Over 3% of new tuberculosis cases exhibit drug resistance, driving an urgent need for new therapies. Multiple studies have highlighted the feasibility of killing *M. tuberculosis* by targeting pathways associated with proteostasis [2–6]; indeed, the ATP-dependent proteases that carry out regulated proteolysis in the mycobacterial cytosol are either strictly essential or conditionally required during infection [7, 8]. This essentiality also makes it challenging to interrogate the physiological roles and regulation of mycobacterial proteases. An expanded understanding of physiological proteolytic pathways would directly support efforts to develop potent and effective protease-targeting anti-mycobacterial therapeutics.

One aspect of the mycobacterial proteolytic landscape that remains poorly characterized is the link between proteostasis and post-translational phosphoarginine (pArg) modifications. Foundational studies in *Bacillus subtilis* found that cytosolic proteins misfolded by proteotoxic stress are labeled with pArg by the arginine kinase McsB [9–12]. pArg modifications are directly recognized by the ATP-dependent unfoldase ClpC through dual binding sites in its small non-catalytic N-terminal domain (NTD), leading to degradation of pArg-marked substrates by the ClpCP protease [11]. Although no arginine kinase has yet been identified in mycobacteria, several pieces of evidence support the existence of an analogous pArg-dependent proteolytic pathway: i) phosphoproteomic studies have detected pArg on *Mycolicibacterium smegmatis* proteins [13, 14]; ii) residues that stabilize pArg binding are strongly conserved in the NTD of the mycobacterial ClpC ortholog ClpC1 [13, 15, 16]; and iii) ClpC1 binds pArg with low micromolar affinity *in vitro* [13, 17]. Nevertheless, the broader regulatory paradigms governing pArg-dependent degradation in mycobacteria remain undefined.

The ClpC1 NTD is also the binding site for several antibiotics that kill mycobacteria by dysregulating the ClpC1P1P2 protease [18–21]. These include the structurally related cyclic peptides cyclomarin A (CymA) and rufomycin (Ruf), as well as the unrelated ecumicin (Ecu). Recent work by Taylor and colleagues demonstrated that CymA also binds to a small non-essential and non-proteolytic mycobacterial protein annotated as ClpC2 [22], which has homology to the ClpC1 NTD but lacks the ATPase modules required for substrate unfolding (**Fig. 1A**). ClpC2 was found to be a transcriptional repressor that binds tightly to an operator sequence overlapping the *clpC2* promoter, thereby suppressing *clpC2* expression under normal growth. Binding of CymA to ClpC2 disrupts its interaction with DNA, which allows transcription and drives strong ClpC2 expression. Accumulated ClpC2 acts as an antibiotic sponge, reducing the amount of CymA available to bind ClpC1, thereby conferring partial protection against CymA toxicity. However, it remains unclear whether ClpC2 functions primarily as an antibiotic defense mechanism, or whether stimuli other than CymA are capable of modulating ClpC2’s repressor activity.

**Figure 1.**
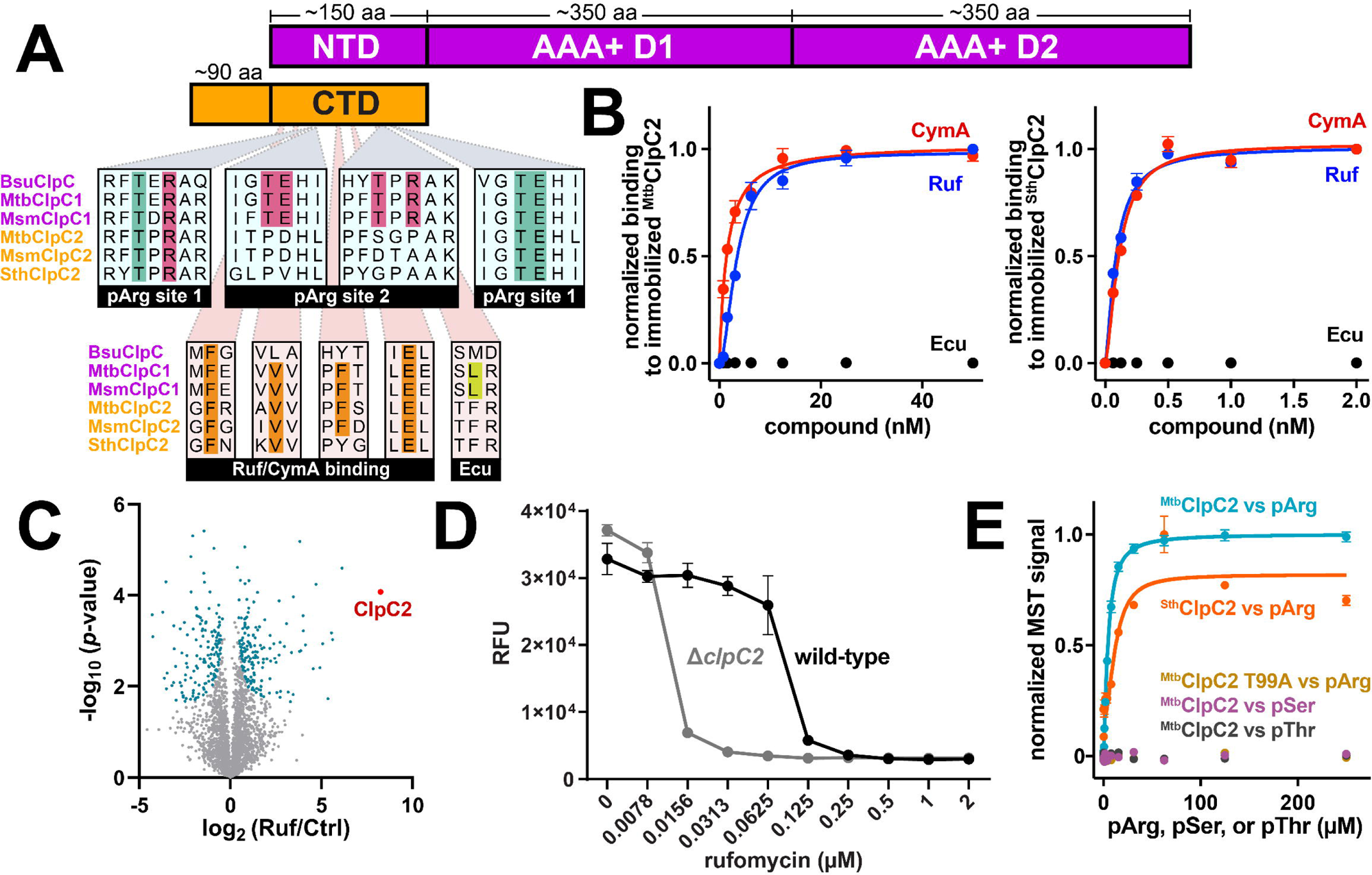
ClpC2 orthologs bind ClpC1-targeting antibiotics and phosphoarginine. A) Domain organization of ClpC1 (purple) and ClpC2 (orange). The ClpC1 NTD and ClpC2 CTD are homologous, sharing ∼40% sequence identity. Sequence alignments show key residues involved in phosphoarginine (pArg) or antibiotic binding. B) Binding of antibiotics to immobilized ClpC2 was measured by biolayer interferometry. Cyclomarin A (CymA) and rufomycin (Ruf) bound to ^Mtb^ClpC2 with *K*_D_ ≈ 1.5 ± 0.1 nM and 3.3 ± 0.3 nM, respectively (left panel) and to ^Sth^ClpC2 with *K_D_*of 86 ± 6 nM and 110 ± 8 nM, respectively (right panel). Ecumicin (Ecu) did not bind to either ortholog. C) Volcano plot illustrating differences in protein abundance in *M. smegmatis* treated with 0.5 µM Ruf, compared to a DMSO-treated control. Differentially enriched proteins that exceeded the FDR threshold of 0.05 are shown in teal. ClpC2 (red point) was the most enriched protein upon Ruf treatment. D) Cultures of wild-type and Δ*clpC2 M. smegmatis* were incubated with varying concentrations of Ruf, and viability was assessed by resazurin fluorescence. E) Binding of ^Mtb^ClpC2 or ^Sth^ClpC2 to pArg, phosphoserine (pSer) or phosphothreonine (pThr) was measured by microscale thermophoresis. pArg bound to ^Mtb^ClpC2 with *K_D_* ≈ 4.7 ± 0.5 µM and to ^Sth^ClpC2 with *K_D_* ≈ 12 ± 0.2 µM. No binding was observed for pSer or pThr to ^Mtb^ClpC2, nor between pArg and a T99A variant of ^Mtb^ClpC2. Binding constants are reported ± standard fit error. Error bars indicate SD among three replicates.

Interestingly, one of the two pArg binding sites present in the ClpC1 NTD is conserved across ClpC2 orthologs, allowing ClpC2 to bind pArg with low micromolar affinity (**Fig. 1A, 1B**) [15]. It has been proposed that ClpC2 serves as a chaperone for pArg-labeled proteins, sequestering them from aggregation and preventing overload of ClpC1P1P2, until the protease can clear them from the cell [15]. We questioned whether binding of pArg may additionally modulate transcriptional repression by ClpC2, analogously to the modulation observed with CymA. Here, we use a suite of biophysical approaches and transcriptional assays to interrogate how pArg and antibiotics perturb ClpC2 oligomerization, DNA binding, and repressor function. Our results reveal that i) ClpC2 oligomerization is required for tight binding to operator DNA, ii) both ClpC1-targeting antibiotics and pArg disrupt ClpC2 oligomerization and DNA binding, and iii) pArg-bearing proteins relieve transcriptional repression by ClpC2. These findings suggest a physiological role for ClpC2 as a pArg-sensitive regulator linking elevated cytosolic pArg levels to downstream transcriptional changes, with a potential role during proteotoxic stress.

## RESULTS

### ClpC2 homologs have conserved binding sites for antibiotics and pArg –

ClpC2 is a small transcriptional repressor present in mycobacteria and various other Actinomycetota species [15]. The C-terminal domain (CTD) of ClpC2 is homologous to the NTD of ClpC1, with ∼40% sequence identity and ∼2.8 Å RMSD between published crystal structures [15, 19, 22]. A prior study by Taylor and colleagues found that the antibiotic cyclomarin A (CymA), which targets the ClpC1 NTD with low nanomolar affinity, also binds to ClpC2 with *K*_D_ ∼2 nM [22]. Indeed, residues important for binding of CymA and the structurally similar antibiotic rufomycin (Ruf) to the ClpC1 NTD are conserved among ClpC2 orthologs (**Fig. 1A**). By contrast, a critical Leu required for binding of the unrelated antibiotic ecumicin (Ecu) is altered to a destabilizing Phe in ClpC2 [23]. We carried out biolayer interferometry (BLI) assays to measure binding of CymA and Ruf to immobilized ^Mtb^ClpC2 (**Table S1**). Our data confirm that CymA binds to ClpC2 with low nanomolar affinity (*K*_D_ ≈ 1.5 nM; **Fig. 1B**) [22], and reveal similarly tight binding for Ruf (*K*_D_ ≈ 3.3 nM; **Fig. 1B**). Notably, both antibiotics also bound to a ClpC2 ortholog from the thermophilic actinobacterium *Streptomyces thermoviolaceus* (^Sth^ClpC2; RefSeq ID: WP_189427858.1; 63% sequence identity to *M. tuberculosis* ClpC2), but with >35-fold higher binding constants (*K*_D_ ≈ 86 nM for CymA, 110 nM for Ruf; **Fig. 1B**). Weaker binding to ^Sth^ClpC2 may be due to the substitution of a critical Phe at position 172 in *M. tuberculosis* to Tyr in *S. thermoviolaceus* (**Fig. S1**): an analogous F80Y substitution in the *M. tuberculosis* ClpC1 NTD weakened antibiotic binding by ∼10-fold [18]. Ecu did not bind to either ortholog, in line with the destabilizing Leu Phe substitution (F184 in both *M. tuberculosis* and *S. thermoviolaceus*).

Taylor and colleagues additionally observed that CymA disrupts ClpC2 binding to its operator sequence within the *clpC2* promoter, leading to strong ClpC2 overexpression [22]. We found that treatment of *M. smegmatis* with Ruf similarly drives ClpC2 overexpression to ∼300-fold of pre-treatment levels (**Fig. 1C**; **Table S2**). Overexpressed ClpC2 is known to sequester CymA, reducing the amount of antibiotic available to bind ClpC1, and thereby diminishing CymA potency [15, 22]. Consistent with this model, we found that deletion of *clpC2* lowers the minimum inhibitory concentration (MIC) of Ruf by ∼8-fold (**Fig. 1D**). The ability of ClpC2 to provide partial protection against CymA and Ruf suggests that ClpC2-mediated sequestration is a common liability for many CymA-like antibiotics.

Sequence analysis also revealed that one of the two phosphoarginine (pArg) binding sites present in the ClpC1 NTD is conserved in the ClpC2 CTD (**Fig. 1A**), which has been noted previously and observed crystallographically [15]. Microscale thermophoresis (MST) confirmed that pArg – but not phosphoserine or phosphothreonine – binds to *M. tuberculosis* ClpC2 (^Mtb^ClpC2) with *K*_D_ ≈ 5 µM (**Fig. 1E**; **Table S1**). Mutation of a conserved Thr in the pArg binding site to Ala (T99A) abolished binding. pArg also bound to ^Sth^ClpC2 (*K*_D_ ≈ 12 µM), suggesting that the ability to bind pArg is a conserved feature among ClpC2 orthologs.

### pArg and rufomycin alter ClpC2 oligomerization –

The molecular mechanism by which antibiotics or other ligands influence ClpC2 DNA binding remains unclear. ClpC2 recognizes a pseudopalindromic operator sequence [22], suggesting that it binds DNA as a dimer. Indeed, Taylor and colleagues showed that ClpC2 forms predominantly dimers by analytical size exclusion chromatography (SEC) [22]. During purification of ^Mtb^ClpC2 and ^Sth^ClpC2 by preparative SEC, we noted that these ∼26 kDa proteins eluted as a mixture of higher-order species (**Fig. 2A**). By contrast, truncated constructs consisting only of the CTD (ClpC2^CTD^) migrated as smaller species, indicating that the N-terminal module is required for higher-order assembly. To better understand the concentration dependence of self-association, we monitored the thermophoretic signal of fluorescently labeled ^Mtb^ClpC2 in the presence of excess unlabeled ^Mtb^ClpC2. The resulting data show concentration-dependent changes in signal, with half-maximal shifts at ∼9 µM (**Fig. 2B**). We further analyzed ^Mtb^ClpC2 by sedimentation velocity analytical ultracentrifugation (SV AUC) at concentrations spanning the self-association curve (**Fig. 2C**). The distribution of ClpC2 oligomers transitions from predominantly dimer at 3.1 µM to predominantly tetramer/octamer at 100 µM. The relatively high *EC_50_*for self-association may explain why higher order oligomers were not observed in prior analytical SEC experiments. The existence of higher order species suggests that multiple self-association interfaces exist on ClpC2, which may be disrupted by ligand binding.

**Figure 2.**
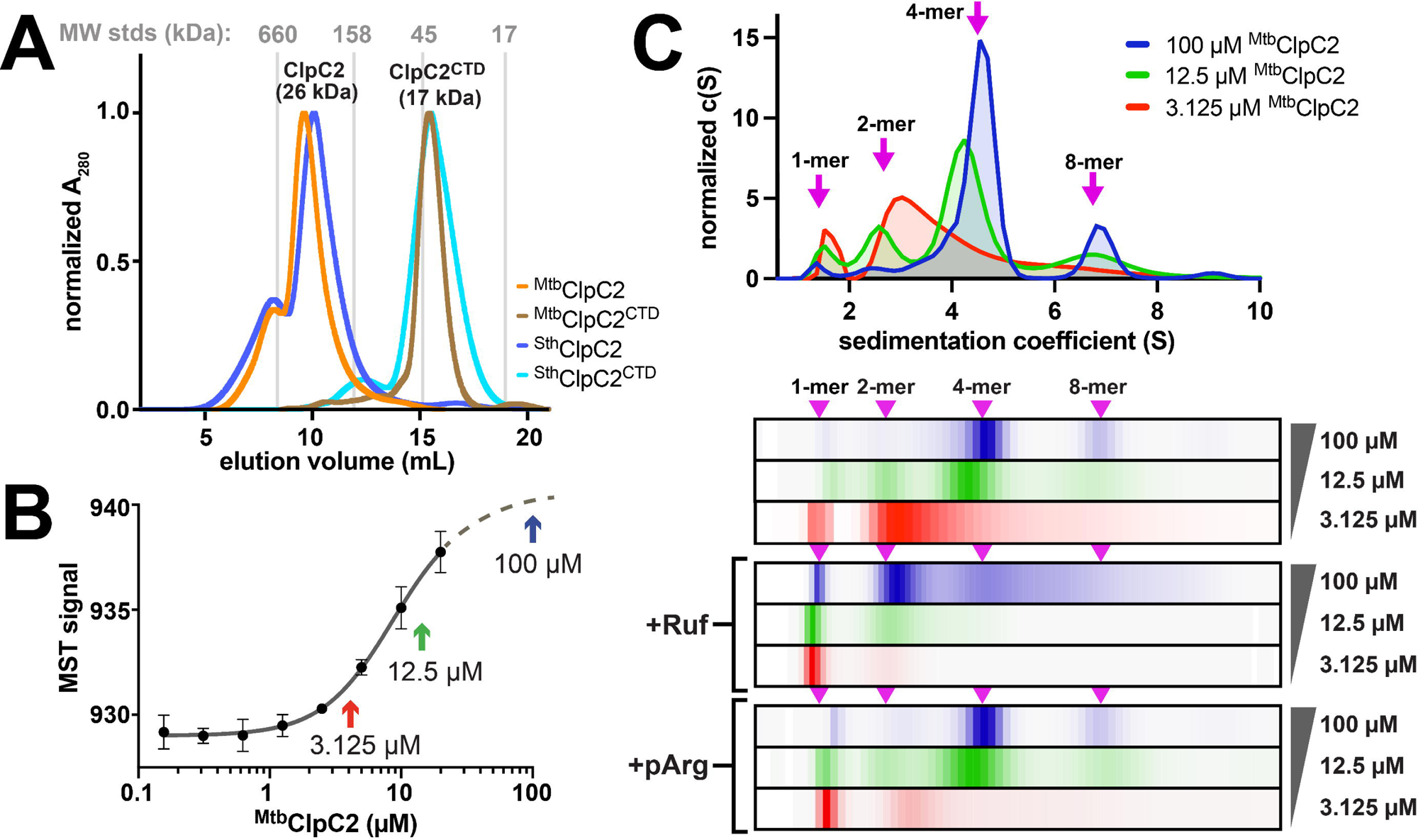
Phosphoarginine and rufomycin disrupt ClpC2 oligomerization. A) Size exclusion chromatograms of full-length or CTD-only constructs of ^Mtb^ClpC2 and ^Sth^ClpC2 show oligomerization. B) The thermophoretic signal from 0.2 µM of fluorescently labeled ^Mtb^ClpC2 shifts upon addition of excess unlabeled ClpC2, suggesting higher-order self-association. Arrows indicate concentrations subsequently tested by ultracentrifugation. Error bars indicate SD among three replicates. C) The indicated concentrations of ^Mtb^ClpC2 were analyzed by velocity analytical ultracentrifugation and fit to sedimentation coefficient distributions shown as plots (upper) and one-dimensional heatmaps (lower) where darker colors correspond to higher *c(S)*. Increasing concentrations of ^Mtb^ClpC2 drive formation of higher order species. Inclusion of 1.2 molar excess of Ruf or 10 molar excess of pArg destabilized oligomers in favor of lower order species. Arrows indicate expected sedimentation coefficients of the indicated species.

To directly assess the consequences of ligand on oligomerization, we examined the effects of saturating Ruf or pArg on ^Mtb^ClpC2 by SV AUC (**Fig. 2C**). At 3.1 µM and 12.5 µM ClpC2, Ruf disrupted oligomers in favor of monomer, while at 100 µM the tetramer/octamer distribution was reduced to monomer/dimer. Phosphoarginine had a similar effect at 3.1 µM ClpC2, favoring monomer formation, and caused modest oligomer disruption at higher ClpC2 concentrations. Thus, both antibiotic and pArg destabilize ClpC2 oligomers.

### pArg releases ClpC2 from its operator –

ClpC2 is known to bind to the *clpC2* operator (*clpC2*o) and thereby repress *clpC2* transcription [22]. We used biolayer interferometry to assess binding of *M. smegmatis* ClpC2 (^Msm^ClpC2) to its cognate *clpC2* operator (^Msm^*clpc2*o), which incorporates two adjacent operator sequences (**Fig. 3A**, **S2A**) [22]. By measuring the equilibrium binding response over a range of ^Msm^ClpC2 concentrations, we determined a *K*_D_ for operator binding of ∼10 nM (**Fig. 3B, 3C**; **Table S3**), in line with prior estimates [22]. At first approximation, BLI binding signal is proportional to the mass of the biomolecules bound to the biosensor [24, 25]. Based on the ratios of immobilized 30.8 kDa dsDNA (0.2 nm) to the binding signal at saturation (∼1.2 nm), we estimate that ^Msm^ClpC2 binds to ^Msm^*clpC2*o in ∼6-fold stoichiometric excess, corresponding to an average of ∼3 molecules of ^Msm^ClpC2 to each operator site. While this may reflect a mixture of oligomeric states, the stoichiometry demonstrates that ClpC2 oligomers larger than dimers can bind to operator DNA.

**Figure 3.**
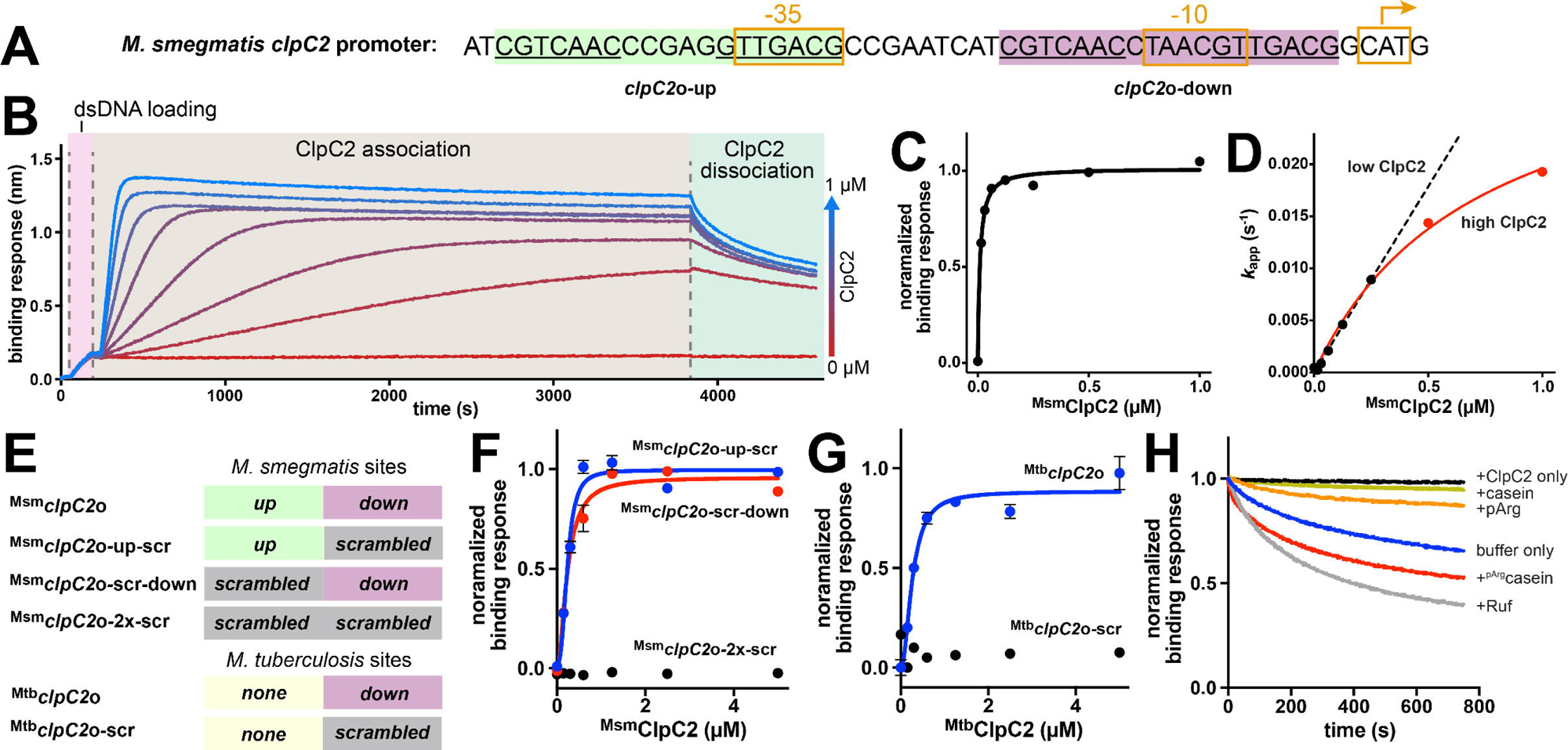
Phosphoarginine and rufomycin disrupt ClpC2 binding to its DNA operator. A) Sequence of the *M. smegmatis clpC2* promoter, with upstream (green) and downstream (purple) ClpC2 operator sites indicated. B) A representative set of biolayer interferometry (BLI) traces shows immobilization of ^Msm^*clpC2o* dsDNA, binding of various concentrations of ^Msm^ClpC2, and dissociation of ^Msm^ClpC2 into buffer. Association and dissociation phases were separately fit to kinetic binding models. C) Equilibrium binding responses of wild-type ^Msm^ClpC2 to ^Msm^*clp*o*C2* were fit to a Hill equation, revealing *K*_D_ of 9.8 ± 2 nM and Hill coefficient of 1.1 ± 0.3. D) Apparent association rates (*k*_app_) were plotted against ^Msm^ClpC2 concentration. At low ^Msm^ClpC2 concentrations, *k*_app_ increased linearly with ^Msm^ClpC2, corresponding an association rate constant (*k*_on_) of 3 10^4^ M^-1^ s^-1^. At higher ClpC2 concentrations *k*_app_ approached a plateau value. E) Representation of *M. smegmatis* and *M. tuberculosis* promoter constructs used for binding studies, with intact or scrambled operator sequences. F) ^Msm^ClpC2 bound to promoter constructs incorporating a single intact operator sequence with *K_D_* ≈ 240 ± 20 nM (^Msm^*clpC2*o-up-scr) or *K_D_* of 250 ± 50 nM (^Msm^*clpC2o*-scr-down). No binding was observed to a construct with both sites scrambled (^Msm^*clpC2*o-2x-scr). G) ^Mtb^ClpC2 bound to its native dsDNA operator (^Mtb^*clpC2*o), containing a single operator sequence, with *K_D_* of 270 ± 20 nM. No binding was observed to a construct with a scrambled operator (^Mtb^*clpC2*o-scr). H) ^Msm^ClpC2 was pre-bound to ^Msm^*clpC2*o, then transferred into wells containing buffer alone, ^Msm^ClpC2 alone, or ^Msm^ClpC2 plus the indicated components. Data points and error bars show mean ± SD of three replicates. Fit values are reported ± standard fit error.

Interestingly, association kinetics were multi-phasic. At high ^Msm^ClpC2 concentrations, binding signal reached a peak and then slowly decreased to a lower equilibrium value (**Fig. 3B**), suggesting that larger oligomers initially bound but dissociated into smaller species. Plotting initial association rates (*k*_app_) revealed a linear relationship between *k*_app_ and protein concentration at lower ^Msm^ClpC2 levels (**Fig. 3D**), corresponding to an association rate constant (*k*_on_) of ∼3×10^4^ M^-1^•s^-1^. However, *k*_app_ values at high ^Msm^ClpC2 concentrations approached a plateau of ∼0.03 s^-1^. This behavior suggests slow interconversion of binding-competent and binding-incompetent states that becomes rate limiting at high concentrations, perhaps reflecting reordering of larger binding-incompetent ^Msm^ClpC2 oligomers into lower-order binding-competent species. Similarly, dissociation rates varied with the amount of bound ^Msm^ClpC2, with higher loading levels leading to faster dissociation (**Fig. S3**).

To assess binding of ^Msm^ClpC2 to a single operator, we constructed variants of ^Msm^*clpC2*o with scrambled sequences in place of the “upstream” (^Msm^*clpC2*o-scr-down), “downstream” (^Msm^*clpC2*o-up-scr), or both operator sequences (^Msm^*clpC2*o-2x-scr) (**Fig. 3E**). ^Msm^ClpC2 bound to ^Msm^*clpC2*o-scr-down or ^Msm^*clpC2*o-up-scr (each incorporating a single operator sequence) with a *K*_D_ of ∼240 nM, while no binding was observed to ^Msm^*clpC2*o-2x-scr (**Fig. 3F**). The fact that ^Msm^ClpC2 binds ∼24-fold tighter to the tandem operator arrangement in wild type ^Msm^*clpC2*o than to variants incorporating a single operator suggests that higher-order ^Msm^ClpC2 complexes bind simultaneously to both sites. While both operator sites are conserved across most *clpC2* promoters, the *M. tuberculosis clpC2* promoter contains only the downstream operator site (**Fig. S2A**) [22]. We tested binding of ^Mtb^ClpC2 to its cognate *clpC2* operator (^Mtb^*clpC2*o) and observed a *K*_D_ of ∼270 nM (**Fig. 3G**), similar to ^Msm^ClpC2 binding to a single site. Given this weaker binding, the *M. tuberculosis clpC2* promoter is likely under less stringent ClpC2-dependent regulation than actinobacterial *clpC2* genes harboring dual operators, and is likely less sensitive to the higher-order oligomerization state of *M. tuberculosis* ClpC2.

We next examined how antibiotics and pArg influence binding of ^Msm^ClpC2 to the promoter region. Biosensors with immobilized ^Msm^*clpC2*o were first incubated with 1 µM ^Msm^ClpC2 to establish a baseline of bound protein, then were transferred to wells containing either ^Msm^ClpC2 alone or ^Msm^ClpC2 together with a stoichiometric excess of ligand (**Fig. 3H**). Binding signal remained stable when transferred to ^Msm^ClpC2 alone. The inclusion of 2 µM Ruf, however, caused rapid dissociation, similar to the effect previously reported with CymA [22]. Inclusion of 500 µM pArg caused partial dissociation of ^Msm^ClpC2, while 10 µM ^pArg^casein (phosphorylated *in vitro* by the *B. stearothermophilus* arginine kinase McsB [11]) triggered more extensive dissociation. The notably greater potency of ^pArg^casein over pArg alone suggests either that ^pArg^casein binding is stabilized by additional contacts to residues on the polypeptide, or that multivalent pArg interactions with *^Msm^*ClpC2 stabilize tighter overall binding. By either model, our results demonstrate the potential for physiological phosphoarginine-bearing proteins to induce ClpC2 dissociation from operator sites.

### Crystal structures of ClpC2 reveal a dimeric interface sensitive to pArg binding –

One mechanism by which ligand may influence oligomerization is direct occlusion of an interaction surface. Crystal structures of the isolated mycobacterial ClpC2 NTD (ClpC2^NTD^) and CTD (ClpC2^CTD^) have been reported, but none shows a direct connection between ligand binding and self-association. *M. tuberculosis* ClpC2^NTD^ crystallized as a dimer [22], but this domain is not known to interact with antibiotic or pArg. Structures of ^Mtb^ClpC2^CTD^ and ^Msm^ClpC2^CTD^ have been determined in complex with CymA [22] and pArg [15], respectively, but the absence of a ligand-free comparator structure makes it difficult to assess the consequences of ligand binding.

To help identify relevant assembly interfaces, we sought to determine the structure of the ClpC2^CTD^ in the absence of interaction partners. For these studies we used ClpC2^CTD^ from the thermophile *S. thermoviolaceus* (^Sth^ClpC2^CTD^; residues 84 - 263), both because thermophilic proteins have a track record of crystallizing well [26] and because it gave us the potential to identify interactions that are conserved across species. Crystals of ^Sth^ClpC2^CTD^ diffracted to 1.41 Å, and the structure was solved by molecular replacement using ^Mtb^ClpC1^NTD^ (PDB: 6PBA) as a search model (**Table S4**). A single molecule of ^Sth^ClpC2^CTD^ occupied the asymmetric unit with residues 93 – 241 resolved (**Fig. 4A**), revealing a similar overall fold to mycobacterial ClpC2^CTD^ (backbone RMSD ∼1.3 Å; **Fig. 4B**). Notably, symmetry-related copies of ^Sth^ClpC2^CTD^ formed a 2-fold-symmetric interface (**Fig. 4A**), stabilized by projection of negatively charged residues (E237, D238) from helix α8 in one protomer into a pocket formed by the N-term of α1, loop α2-α3 and loop α5-α6 of the second protomer, mutually burying 2,140 Å^2^. Residues that line the pocket are conserved across ClpC2 orthologs (**Fig. S1**). By contrast, there is lower sequence conservation at the end of helix α8 and in the C-terminal tail, which varies in length. However, Jpred secondary structure prediction of orthologs consistently predicts that 8 ends near a conserved hydrophobic residue (L235 in *M. tuberculosis* and *M. smegmatis;* L236 in *S. thermoviolaceus*) and is followed by a polar or negatively charged residue (T236 in *M. tuberculosis*; E236 *M. smegmatis*; E236 in *S. thermoviolaceus*) (**Fig. S1**), which would allow equivalent contacts within the pocket. A similar but less intimate interaction occurs between symmetry related chains in the reported structure of ^Mtb^ClpC2^CTD^ bound to CymA (**Fig. 4C**) [22], with projection of 8 into the pocket partially blocked by a bound sulfate ion. Together, these observations suggest that this interface is evolutionarily conserved.

**Figure 4.**
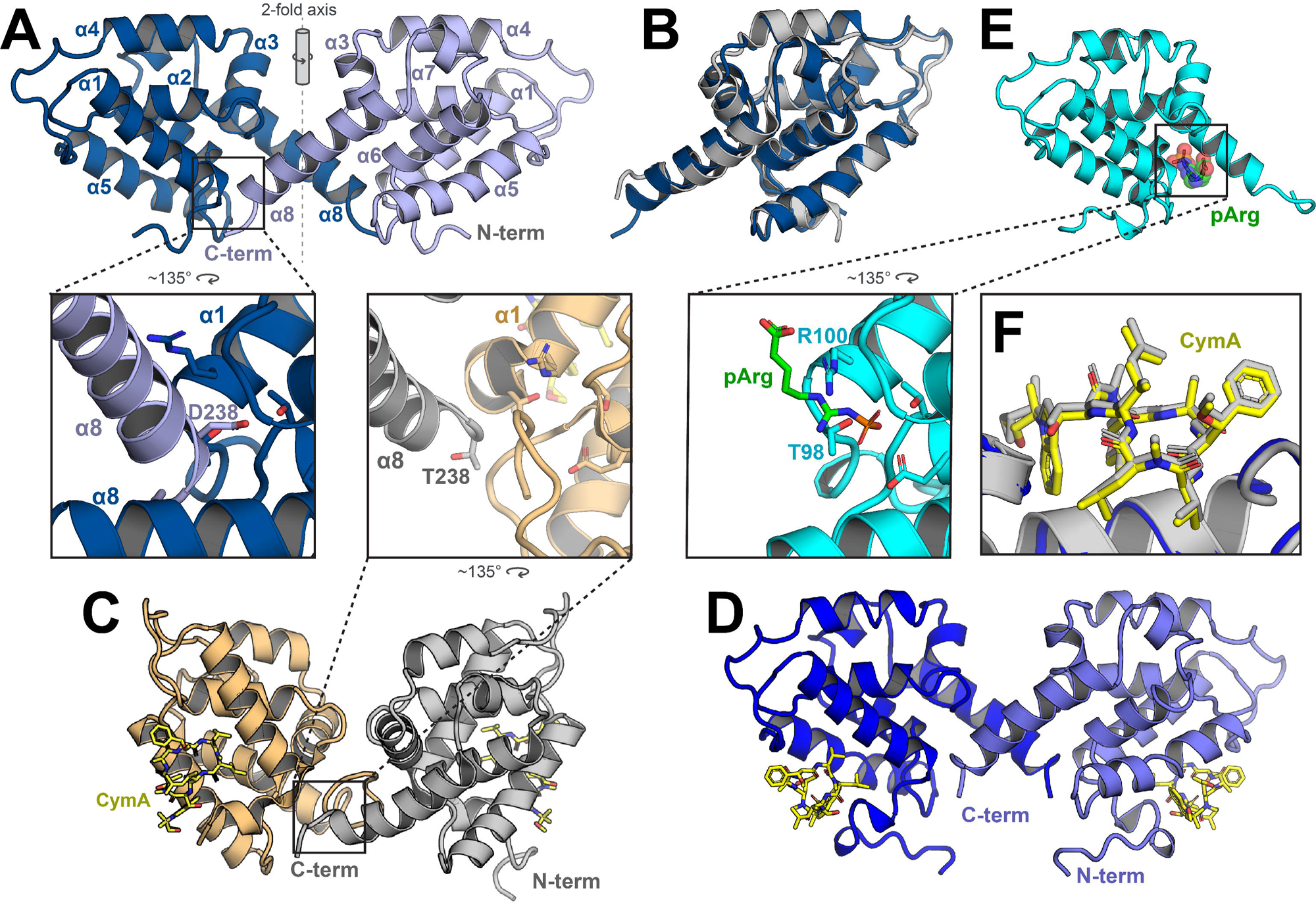
Phosphoarginine binding sterically blocks a dimerization interface in the ClpC2 C-terminal domain. **A**) The crystal structure of *S. thermoviolaceus* ClpC2 CTD reveals a dimer formed from subunits in two asymmetric units that cross a crystallographic 2-fold axis. The dimer interface is stabilized by the C-terminal helix, including Asp238. **B**) An overlay of *S. thermoviolaceus* ClpC2 CTD (blue) and *M. tuberculosis* ClpC2 CTD (gray; PDB: 8AD9) shows close structural homology (1.3 Å RMSD). **C**) The previously reported crystal structure of *M. smegmatis* ClpC2 CTD in complex with CymA (PDB: 8B9O) reveals a similar dimer arrangement as in A, also stabilized by the C-terminal helix. **D**) The crystal structure of *S. thermoviolaceus* ClpC2 in complex with CymA (yellow) shows an identical dimer arrangement as in A (0.4 Å RMSD. **E**) The co-crystal structure of *S. thermoviolaceus* ClpC2 (cyan cartoon) in complex with pArg (transparent spheres) reveals that pArg prevents dimer formation by sterically blocking projection of the C-terminal helix into its binding pocket. **F**) Structural alignment of *S. thermoviolaceus* ClpC2 (gray) or *M. smegmatis* ClpC2 CTD (blue) bound to CymA shows identical binding modes.

To test the effects of CymA on the *^Sth^*ClpC2^CTD^ dimeric interface, we soaked *^Sth^*ClpC2^CTD^ crystals in 1 mM CymA and determined the *^Sth^*ClpC2^CTD^•CymA co-crystal structure to 1.7 Å (**Table S4**). A well-occupied CymA molecule was found at the expected binding site (**Fig. 4D**; **S4A**). CymA did not alter the overall fold (RMSD 0.37 Å) or packing interactions, including the α8-mediated interface. Thus, the reduction in oligomeric state induced by CymA-like antibiotics (**Fig. 2C**) likely arises from effects outside of the α8 interface. (For example, CymA may sterically reposition the NTD in a way that disfavors oligomer assembly.)

Importantly, the conserved pocket into which α8 docks coincides with the expected pArg binding site [15], suggesting that these interactions are mutually competitive. Indeed, the side chain of *S. thermoviolaceus* E237 (T236 in *M. tuberculosis*; E236 in *M. smegmatis*) projects outward from α8 and occupies the same position as pArg in the reported *^Msm^*ClpC2^CTD^•pArg structure [15], making polar contacts with T103 and T200 (T100 and T197 in *M. tuberculosis* ClpC2). Soaking ligand-free ^Sth^ClpC2^CTD^ crystals in 50 µM pArg caused rapid cracking and disintegration; however, we were able to obtain a 1.3 Å structure of ^Sth^ClpC2^CTD^ bound to pArg by co-crystallization (**Table S4**). *^Sth^*ClpC2 bound pArg in a similar manner to that observed in the reported mycobacterial ClpC2•pArg structure [15], with the phospho-guanidinium group well resolved in the electron density map (**Fig. 4E**; **S4B**). The presence of pArg only modestly altered the overall conformation of ^Sth^ClpC2^CTD^ (RMSD 0.77 Å), but ^Sth^ClpC2^CTD^•pArg crystals notably adopted a different packing arrangement without an α8-mediated interface, consistent with steric hindrance by pArg. These structural details agree with our SV AUC data (**Fig. 2C**) and suggest that, in contrast to CymA or Ruf, modulation of quaternary state by pArg arises, at least in part, through disruption of the crystallographically observed ClpC2 CTD dimerization interface.

To further probe whether the α8-mediated interface stabilizes higher order ClpC2 oligomerization, we mutated *M. tuberculosis* T236 to a bulkier Lys, which would be expected to sterically block the interface noted in the ^Mtb^ClpC2^CTD^ structure and electrostatically repel R100. By SV-AUC, we observe that ^Mtb^ClpC2^T236K^ adopts lower-order species than wild-type ^Mtb^ClpC2 at 12.5 µM and 3.1 µM (**Fig. S5A**), confirming that disruption of the 8/pocket interaction destabilizes higher-order oligomerization. Additionally, Taylor and colleagues report a mutation in the ClpC2 NTD, R56A, that removes a salt bridge stabilizing NTD dimers and causes ^Mtb^ClpC2 to adopt predominantly monomers by SEC. We analyzed ^Mtb^ClpC2^R56A^ by SV-AUC and saw a similar shift at low concentrations (**Fig. S5B**). Together our results suggest that higher order ClpC2 oligomerization is stabilized by multiple self-association interfaces.

### pArg relieves transcriptional repression by ClpC2 –

To further interrogate how pArg modulates ClpC2 activity, we used a fluorescent *in vitro* assay to test transcriptional repression by ^Mtb^ClpC2 in the presence of different ligands. We constructed a DNA template incorporating a consensus *Escherichia coli* σ70 promoter [27] followed by the *M. tuberculosis clpC2* operator (^Mtb^*clpC2*o; **Fig. S2B**) [22] and a sequence encoding the F30-Broccoli aptamer [28] (**Fig. 5A**). F30-Broccoli binds the non-fluorescent ligand DFHBI-1T to produce a complex with GFP-like fluorescence [28, 29], allowing us to follow fluorescence as a readout for transcriptional activity. In the presence of *E. coli* RNA polymerase and σ70, we observed an increase in fluorescence over time, demonstrating a baseline level of transcriptional activity (**Fig. 5B**). Inclusion of 6 µM ^Mtb^ClpC2 suppressed the initial rate of transcription to 25%. ^Mtb^ClpC2 did not repress transcription of a template with a scrambled operator sequence (**Fig. 5C**). Additionally, the RNA polymerase inhibitor rifampin [30] halted all transcriptional activity (**Fig. 5C**). No repression was observed by ^Mtb^ClpC2 constructs consisting of NTD or CTD alone (**Fig. S6**). Next, we titrated ^Mtb^ClpC2 and found that the initial transcriptional rate was inhibited with an *IC*_50_ of ∼1.4 µM (**Fig. 5D**). This confirms the ability of ^Mtb^ClpC2 to repress transcriptional activity when bound to its cognate operator.

**Figure 5.**
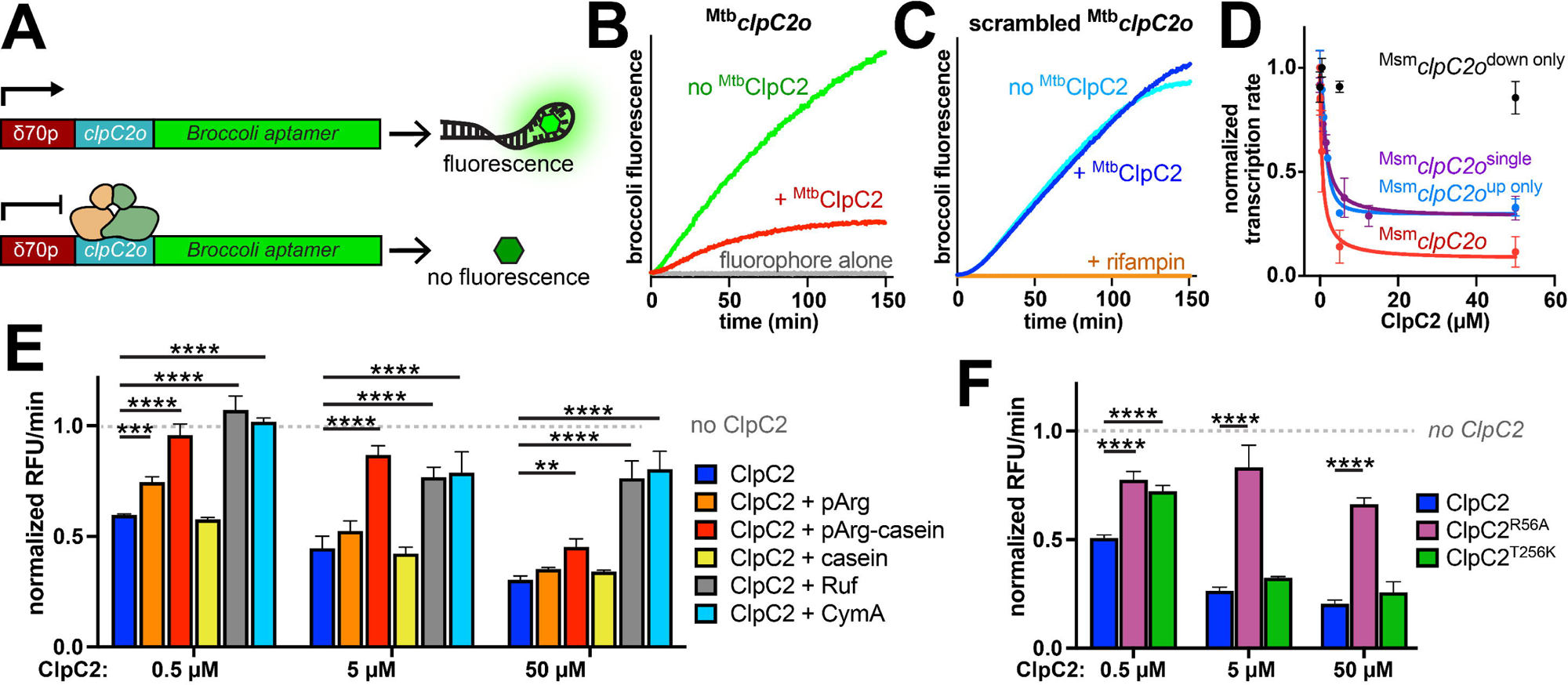
Phosphoarginine and antibiotics relieve transcriptional repression by ClpC2 *in vitro*. A) Diagram of the DNA template used for *in vitro* transcription studies, including a σ70-dependent *E. coli* promoter, *clpC2* operator, and sequence encoding a F30-broccoli RNA aptamer. B) Transcription of the broccoli aptamer was monitored by the increase in fluorescence over time (green trace). Inclusion of 6 µM ^Mtb^ClpC2 decreased transcription (red trace). DFHBI-1T alone produced minimal fluorescence (gray trace). C) Transcription from a template incorporating a scrambled operator sequence was similar in the absence (cyan) or presence (blue) of 6 µM ^Mtb^ClpC2. By contrast, 2 µM rifampin completely inhibited transcription (orange trace). D) Transcriptional rate was measured as a function of ^Msm^ClpC2 concentration, using templates with scrambled or intact operator sequences. E) Transcription was measured in the presence of the indicated concentration of ^Mtb^ClpC2, or with the additional inclusion of 500 µM pArg, 10 µM ^pArg^casein, 10 µM unphosphorylated casein, 1.2-fold molar excess Ruf or 1.2-fold molar excess CymA. F). Transcriptional repression is shown for the indicated concentration of wild-type ^Mtb^ClpC2, and for ^Mtb^ClpC2 variants incorporating R56A or T256A mutations. Error bars reflect standard deviation of ≥3 replicates. Statistical significance was determined by ordinary one-way ANOVA followed by Dunnett’s multiple comparison test to compare each condition to ClpC2 alone (blue bar) at a given concentration. Multiplicity-adjusted *p*-values are indicated as follows: * *p* ≤ 0.05, ** *p* ≤ 0.01, *** *p* ≤ 0.001, **** *p* ≤ 0.0001. Non-significant comparisons are not shown.

To investigate the effects of dual operators on repression, we evaluated transcriptional repression by ^Msm^ClpC2 on cassettes incorporating ^Msm^*clpC2*o (**Fig. S2B**). We measured an *IC*_50_ of 630 nM and a maximum reduction of the initial transcriptional rate to ∼10% (**Fig. 5D**). Weaker *^Msm^*ClpC2-dependent repression was measured for a cassette incorporating ^Msm^*clpC2*o-up-scr, with *IC*_50_ ≈ 1.5 µM (**Fig. 5D**). The tighter repression of the dual-operator template supports a model in which ClpC2 oligomers simultaneously interact with adjacent operator sequences to achieve stronger transcriptional repression. Negligible repression was observed for a template incorporating ^Msm^*clpC2*o-scr-down (**Fig. 5D**), likely because the positioning of the single downstream operator was too distant to block RNA polymerase binding, suggesting that bound ClpC2 represses transcription by preventing the initial engagement of RNA polymerase with RNA rather than by blocking elongation.

To test the effects of ClpC1-targeting antibiotics and pArg on repression, we carried out *in vitro* transcription assays with 0.5 µM, 5 µM, or 50 µM ^Mtb^ClpC2 in the absence or presence of CymA (1.2-fold molar excess), Ruf (1.2-fold molar excess), pArg (500 µM), ^pArg^casein (10 µM), or unmodified casein (10 µM) (**Fig. 5E**). As expected, based on the ability of Ruf and CymA to drive ClpC2 overexpression (**Fig. 1C**), these antibiotics strongly counteracted ^Mtb^ClpC2 repression, increasing transcription >50% in the context of 50 µM ^Mtb^ClpC2 and fully restoring baseline activity in the presence of 5 µM or 0.5 µM ^Mtb^ClpC2. pArg alone only modestly elevated transcription, with a ∼5% increase at the lowest ^Mtb^ClpC2 concentration. Strikingly, ^pArg^casein had a much stronger effect, restoring transcription to near baseline levels with 0.5 µM or 5 µM ^Mtb^ClpC2. Unmodified casein did not alter ^Mtb^ClpC2 repression.

To confirm that repression correlates with ClpC2’s oligomeric state, we tested the ability of full-length ^Mtb^ClpC2 bearing either T236K or R56A mutations to repress transcription (**Fig. 5F**). At 0.5 µM ^Mtb^ClpC2, the T236K mutation partially impaired repression. R56A exhibited strongly impaired transcription at all concentrations tested. Thus, mutations that disrupt ClpC2 oligomerization correspondingly impair transcriptional repression. Together, these results demonstrate that pArg-modified proteins relieve transcriptional repression by ClpC2, providing a physiological mechanism by which elevated proteomic pArg can activate downstream transcriptional programs.

## DISCUSSION

Phosphoarginine is one of few post-translational degrons known to exist in bacteria, and its specific role in mycobacterial physiology remains an intriguing mystery. Evidence suggests that pArg-bearing proteins are recognized as proteolytic substrates by mycobacterial ClpC1P1P2 [13, 15, 17], but the broader regulatory landscape surrounding pArg modifications is unknown. Here, we show that ClpC2 can mechanistically link pArg levels to transcriptional regulation of target genes. Taylor and colleagues previously found that ClpC2 functions as a transcriptional repressor, and that binding of the ClpC1-targeting antibiotic CymA to ClpC2 relieves negative-feedback repression of the *clpC2* gene [22]. We observed similar behavior here with Ruf, a structurally related antibiotic. Additionally, building on reports that ClpC2 binds pArg [15], we show that pArg modulates the oligomeric state, DNA binding ability, and repressor activity of ClpC2. Our findings suggest that ClpC2 mechanistically links the abundance of pArg modifications to the downstream transcriptional regulation of *clpC2*, and possibly other mycobacterial genes.

The elevated expression of ClpC2 induced by pArg may allow it to sequester pArg-bearing substrates and thereby prevent them from interfering with important ClpC1P1P2-dependent proteolytic pathways. Indeed, sequestration of ClpC1 substrates by ClpC2 has been proposed previously and demonstrated *in vitro* [15]. However, our data indicate that only some classes of substrates stimulate ClpC2 overexpression: ^pArg^casein drove dissociation of ClpC2 from operator DNA, while unmodified casein did not. This suggests a mechanistic distinction between general proteotoxic stress and specific pArg-linked stress conditions. Our data also suggest that pArg-sensing is a broadly conserved function of ClpC2 across Actinomycetota. While it is unclear whether CymA-like antibiotics are common enough in nature for antibiotic resistance to be the primary physiological role of ClpC2, the strict conservation of the ClpC NTD pArg binding modules across Actinomycetota [13, 16] implies that pArg-dependent pathways are widely distributed.

The biophysical and functional studies here highlight the importance of ClpC2 oligomerization in DNA binding. The *clpC2* operator sequence is pseudopalindromic, implicating ClpC2 dimers as the fundamental DNA binding unit [22]. Notably, *M. smegmatis* and most other mycobacteria possess two adjacent operators in the *clpC2* promoter region [22], potentially allowing multivalent engagement by higher order ClpC2 assemblies. Indeed, we find that higher-order ClpC2 oligomers bind substantially tighter to double than to single operator sequences, and that ligand-induced reduction in oligomeric state drives dissociation of ClpC2 from DNA. Multiple lines of evidence now suggest that ClpC2 assemblies are stabilized by multiple interfaces, providing multiple opportunities for ligand binding to destabilize oligomer formation: our studies identify a specific pArg-sensitive interface within the ClpC2 CTD; ClpC1-targeting antibiotics destabilize oligomers through a distinct CTD binding site; and prior reports identified a mutation in the ClpC2 NTD that also prevents higher order assembly [22]. Further structural studies will be required to fully understand the interplay between ClpC2 oligomerization and DNA binding.

Our experiments also predict regulatory differences between *clpC2* promoters containing double operator sites and those incorporating a single, which binds ClpC2 with ∼25-fold weaker affinity. In *M. smegmatis* and other mycobacteria with double operators [22], low cytosolic levels of ClpC2 are likely sufficient to suppress *clpC2* transcription. Thus, high levels of pArg modifications (perhaps correlated with strong proteotoxic stress, such as heat shock [15]) may be required to fully dissociate ClpC2 from the *clpC2* promoter and facilitate high level expression. By contract, the single operator in *M. tuberculosis* may allow a greater dynamic range of regulation, such that lower pArg levels can effectuate physiologically meaningful changes in repressor binding and *clpC2* expression. By contrast, ClpC1-targeting antibiotics that bind ClpC2 with low-nanomolar affinity may effectively relieve repression in both systems. (Interestingly, a recent study found that the cyclomarin variant desoxycyclomarin drove ClpC2 overexpression in *M. tuberculosis*, while the structurally related ilamycin E did not [31]. This difference may arise from the ∼25-fold weaker binding of ilamycin E to ClpC2.)

Taken together, our findings position mycobacterial ClpC2 as a pArg-responsive transcriptional regulator, able to sense pArg levels and accordingly modulate downstream gene expression. The details of this regulatory phenomenon may vary across Actinomycetota, depending on the distribution and details of ClpC2 operator sequences. Future studies will be required to unravel the upstream regulation of pArg installation clarify the full spectrum of ClpC2’s physiological roles.

## MATERIALS AND METHODS

### Plasmid construction *–*

Sequences encoding *M. tuberculosis* ClpC2 and *S. thermoviolaceus* ClpC2 were ordered from Twist Biosciences with codon optimization for *E. coli* expression. McsB was amplified from *Geobacillus stearothermophilus* (ATCC 12980) genomic DNA. ^Mtb^ClpC2^CTD^ (86-252) and MecA were each cloned in-frame with an N-terminal 6xHis tag and TEV cleavage site (TEVcs) [32] (MGSSHHHHHHDYDIPTTENLYFQG) in a pET-21a derived plasmid (EMD Millipore). ^Sth^ClpC2^CTD^ was cloned in frame with a C-terminal *Vibrio cholerae* MARTX toxin cysteine protease domain (CPD tag) [33] and 6xHis in pET-22b (EMD Millipore). Additional ^Sth^ClpC2 and ^Mtb^ClpC2 constructs for BLI experiments were cloned in frame with a C-terminal sortase tag (LPETGGHHHHHHH) [34] in a pET-22b vector. ^Gst^McsB was cloned in frame with a C-terminal 6xHis tag in a pET-22b derived plasmid. Mutants were generated by primer-based site-directed mutagenesis. *E. coli* RNAP (pVS10) and [35] and *E. coli* 6xHis-σ70 (pET28-derived) [36] were a kind gift from Dr. Jeremy Bird (University of Delaware). Construct sequences were verified by Sanger (Genewiz) or whole-plasmid sequencing (Plasmidsaurus). Plasmids used in this study are listed in **Table S5**.

### Protein expression and purification –

*E. coli* σ70 and RNAP were expressed and purified as described [35, 36]. All other constructs were transformed into *E. coli* strain ER2566 (NEB) by electroporation and grown in a 1:1 mixture of LB and 2xYT at 37°C to an OD_600_ of 0.8. Overexpression was induced by the addition of isopropyl β-D-1-thiogalactopyranoside (IPTG), and incubation continued for 18 h at 18°C. Cells were harvested by centrifugation, resuspended in Lysis Buffer (25 mM HEPES, 300 mM NaCl, 10 mM imidazole, 10% glycerol, pH 7.5), and lysed by sonication. TEVcs constructs were purified by Ni-NTA (MCLab) chromatography, after which the 6xHis-TEV tag was cleaved by 4°C overnight incubation with 1 µg TEV protease per 50 µg target protein, followed by a second round of Ni-NTA chromatography to remove the tag and TEV protease. ^Sth^ClpC^CTD^ was bound to Ni-NTA resin and self-cleavage of the CPD-6xHis tag was induced by the addition of inositol hexakisphosphate; cleaved ^Sth^ClpC^CTD^ was recovered by elution in Lysis Buffer. McsB was purified by Ni-NTA and anion exchange (Source 15Q, Cytiva) chromatography. All constructs were purified by size exclusion chromatography (Superdex 200 pg 16/600, Cytiva) in CPD buffer (25 mM HEPES, pH 7.5, 200 mM KCl, 10 mM MgCl_2_, 0.1 mM EDTA). Protein concentrations were measured by A_280_. Analytical size exclusion (Superdex 200 Increase 10/300, Cytiva) was performed in CPD buffer with 70 µM injections of ClpC2.

### Microscale thermophoresis –

His-tagged ClpC2 constructs were diluted to 100 nM and incubated with RED-tris-NTA (NanoTemper) or HIS Lite OG488-Tris NTA-Ni dye (AAT Bioquest) in CPD buffer with 0.05% w/v Tween 20 for 45 min at room temperature. Phosphoarginine (Toronto Research Chemicals), phosphoserine (MedChemExpress), and phosphothreonine (MedChemExpress) were serially diluted in CPD + 0.05% Tween 20, mixed 1:1 with ClpC2 and dye, and loaded into capillary tubes. Thermophoresis measurements were collected on a NanoTemper Monolith NT.115 instrument at 40% LED and laser power at 25°C. Thermal shift data were fit to a quadratic single site binding equation in Prism (GraphPad).

### Analytical ultracentrifugation –

ClpC2 was mixed with ligand in CPD buffer and allowed to equilibrate at room temperature for 2 h. Samples were loaded into AUC cells, placed into an An-50 Ti 8-hole rotor, and spun at 40,000 RPM in a ProteomeLab XL-I (Beckman-Coulter) analytical ultracentrifuge. Sedimentation velocity scans were collected using interference optics at 5 s intervals. Continuous distribution (*c*(*s*)) analysis was performed in SedFit [37] using buffer density, buffer viscosity, and protein partial specific volume values estimated from SEDNTERP [38].

### X-ray crystallography –

^Sth^Clp2^CTD^ was desalted into a low salt buffer (25 mM HEPES, 50 mM NaCl, pH 7.5) over Sephadex G-10 resin (Cytiva). Crystals were produced by hanging drop vapor diffusion at 20°C in 1.4 µL drops containing equal volumes of protein and crystallization buffer, suspended over a 500 µL reservoir of crystallization buffer. Rectangular crystals of ^Sth^Clp2^CTD^ alone formed from 7.5 mg/mL protein and 0.1 M Bis-tris, pH 5.75, 29% PEG3350 in ∼7 days. Co-crystals with cyclomarin A (CymA; a kind gift from Dr. Scott Franzblau at the Institute for Tuberculosis Research, UIC) were generated by soaking crystals of ^Sth^Clp2^CTD^ in crystallization buffer with 1 mM CymA. Oval-shaped crystals of ^Sth^Clp2^CTD^ in complex with pArg grew from 15 mg/mL protein in 0.1 M Bis-tris, pH 5.5, 1 mM pArg, and 2 M ammonium sulfate over ∼14 days, and were cryoprotected by brief incubation in 0.1 M Bis-tris, pH 5.5, 2.7 M ammonium sulfate, 10% glycerol, and 1 mM pArg.

X-ray diffraction datasets were collected at NSLS-II beamline FMX for ^Sth^Clp2^CTD^ alone and ^Sth^Clp2^CTD^•CymA, and at CHESS beamline 7B2 for ^Sth^Clp2^CTD^•pArg; both at 100 K. Raw data was processed by HKL2000 [39]. The ^Sth^Clp2^CTD^ structure was solved by molecular replacement in Phaser [40] using the structure of the *M. tuberculosis* ClpC1 N-terminal domain [20] (PDB: 6PBA) as a search model. Structures in complex with pArg and CymA were solved using ^Sth^Clp2^CTD^ as search model. Models were built in Coot [41] and refined in Phenix [42]. Structures were deposited into the Protein Database with IDs: 9P2P (^Sth^Clp2^CTD^ alone), 9P39 (^Sth^Clp2^CTD^•pArg), 9P4H (^Sth^Clp2^CTD^•CymA). Surface area calculations of the ClpC2 dimer were performed by PISA PDB [43] and represent total buried volume.

### Biolayer interferometry –

Biotinylated dsDNA was generated either by annealing single stranded DNA oligos (IDT) or by amplifying regions from constructs generated for *in vitro* translation studies (**Fig. S2**). For annealed DNA, single stranded oligos (one of which incorporated a 5’ biotin tag) were reconstituted in 10 mM Tris, pH 8.0, and heated at 95°C in a heat block for 2 min. The heat block was turned off, wrapped in aluminum foil, and allowed to cool to RT for 1 h. For amplified constructs, DNA was PCR amplified using Taq polymerase (Apex) with a 5’ biotin incorporated on the upstream primer. For immobilization of ^Sth^ClpC2 and ^Mtb^ClpC2, purified protein was biotinylated using sortase [34]. 300 µM ClpC2, 10 µM sortase and 1 mM Gly-Gly-Gly-biotin (VectorLabs) were incubated in CPD at 30°C for 2 h and desalted into CPD + 0.05% Tween 20 to remove excess biotin.

Biolayer interferometry (BLI) experiments were carried out on the Octet RH16 (Sartorius) at 25°C in binding buffer (CPD buffer + 0.05% Tween 20; studies with Cym or Ruf additionally included 3% DMSO, which had little effect on binding or dissociation). Biotinylated dsDNA or protein was loaded onto streptavidin coated tips (Sartorius) to a response signal of 0.5 - 1 nm. Initial baselines were established in binding buffer alone. Tips were transferred into wells containing ^Mtb^ClpC2 and/or antibiotic to measure association. Dissociation was monitored by transferring tips to wells with binding buffer alone or binding buffer supplemented with pCasein (1 µM), pArg (500 µM), or Ruf (1.2 µM). Association rates were calculated in Prism by fitting to single- or double-exponential binding models. Initial velocities from the fit were used to determine *k*_on_ rates. Dissociation rates were established by fitting data to a single-exponential model and plotted against ClpC2 or compound concentration.

### *In vitro* transcription assays (IVT) –

*In vitro* transcription (IVT) template constructs were initially cloned into a pET22b-derived plasmid by restriction enzyme cloning using BamHI/HindIII/XhoI sites flanking σ*70p*/*clpC2o*/broccoli aptamer components (IDT), respectively. Template amplicons extending from the 5’ end of the σ70 promoter to the 3’ end of the broccoli aptamer were generated by PCR and purified by spin column. To form the *E. coli* RNAP holoenzyme, 1 µM RNAP and 5 µM σ70 were incubated on ice for 10 min in IVT buffer (40 mM HEPES, 100 mM KCl, 50 mM MgCl_2_, 1 mM DTT, 10% w/v glycerol, pH 7.9). In a separate tube, 200 nM of template DNA was incubated in buffer with or without ClpC2, ^pArg^casein (10 µM), casein (10 µM), pArg (500 µM), Ruf (1.2-fold molar excess), or CymA (1.2-fold molar excess) at 37°C for 15 min. DNA mixtures and holoenzyme were mixed and incubated at 37°C for 15 min, followed by addition of 100 nM of DFHBI-1T (MedChemExpress). To normalize DMSO carryover from antibiotic and fluorophore, all samples were adjusted to a final concentration of 3% DMSO. Reactions were initiated with 5 mM of each NTP. Aptamer transcription was monitored in black non-binding 385-well plates (Grainger) in a Tecan Spark plate reader by following 512 nm fluorescence upon 465 nm excitation.

### Casein arginine phosphorylation –

To generate pArg-bearing casein, 10 µM β-casein (Sigma), 3 µM *Geobacillus stearothermophilus* McsB, and an ATP regeneration system (16 mM creatine phosphate and 0.32 mg/mL creatine kinase) were incubated in CPD buffer supplemented with 1 mM EDTA, 1 mM DTT, and 5 mM ATP for 24 h at 50°C. Casein was purified by size exclusion chromatography (Superdex 200; Cytiva) and the fractions containing pure casein were concentrated.

### Mass spectroscopy –

*Mycolicibacterium smegmatis* MC^2^ 155 cells containing a kanamycin-resistant pNIT-derived vector [44] as a selectable marker were grown in Middlebrook media (HiMedia) at 37°C to an OD_600_ of 0.8 in the presence of 20 µg/ml kanamycin. Cultures were grown to a final OD_600_ ≈ 0.1 and treated with 0.4 µM of Ruf or mock treated with DMSO alone, in triplicate for each condition. Growth continued to an OD_600_ ≈ 0.6, and cells were harvested by centrifugation at 4000 rcf for 30 min at 4°C and resuspended in cold PBS. 1 mL of cells were lysed by bead disruption and clarified by centrifugation at 15,000 rcf for 30 min at 4°C. Protein concentration was normalized by Bradford assay (Bio-Rad).

Samples were prepared for mass spectrometry by filter-aided sample preparation (FASP) [45]. Samples were incubated for 50 min at 56°C in 200 mM DTT, then diluted to 10 mL in 0.1 M Tris, 7.5, 8 M urea, 25 mM 2-iodoacetamide and incubated in the dark at RT for 45 min to alkylate cysteines. Samples were washed twice with 5 mL of Wash 1 (0.1 M Tris, 7.5, 8 M urea) and twice with 5 mL of Wash 2 (0.1 M Tris, 7.5, 4 M urea) at 4000 rcf in a 10,000 MWCO spin concentrator (Millipore Amicon). Samples were buffer exchanged into 100 mM ammonium bicarbonate and concentrated to 1 mL. Proteins were digested with trypsin (Thermo) at a 2000:1 protein:trypsin ratio in 100 mM ammonium bicarbonate overnight at 37°C in a spin concentrator. Peptides were recovered by centrifugation for 10 min at 4000 rcf and further eluted with 0.5 M NaCl. After lyophilization, samples were reconstituted in 0.1% TFA and desalted using a HyperSep C_18_ column.

LC-MS/MS analysis was performed using an Orbitrap Eclipse MS (Thermo Scientific) coupled with an Ultimate 3000 nanoLC system and a FAIMS Pro Interface (Thermo Scientific). Peptides were loaded onto a trap column and then separated by an analytical column (PepMap C18, 3.0 µm; 25 cm x 75 mm I.D.; Thermo Scientific) at 300 nL/min flow rate using a 165-min gradient from Buffer A (0.1% formic acid in water) to Buffer B (0.1% formic acid in acetonitrile). Protein quantitation was performed using the MaxQuant software (version 1.6.4.5) with default parameters, including trypsin as an enzyme with a maximum of two missed cleavage sites; acetylation (protein N-terminal and Lys), and oxidation (Met) as variable modifications; cysteine carbamidomethylation as a fixed modification; minimum peptide length of 7 amino acids; and false discovery rate (FDR) were set at 1% for both protein and peptide identification.

## ACKNOWLEDGEMENTS

The authors thank Vijay Parashar for use of thermophoretic instrumentation; Jeremy Bird for advice on IVT assays; Yanbao Yu for help with mass spectrometry; Farzaneh Hojjati and Alexzandria Sheppard for comments on the manuscript; and members of the Schmitz lab for helpful comments and advice. K.R.S. was supported by award P20GM104316 from NIH NIGMS and award R01AI171196 from NIH NIAID and NIGMS. This work was also supported by a core access award through the Delaware INBRE program, funded by NIH NIGMS P20GM103446 and the State of Delaware. X-ray crystallography resources at the University of Delaware were supported by NIH NIGMS award S10OD26896. The Octet RH16 was supported by NSF MRI 2215833. The University of Delaware Mass Spectrometry Core Facility was additionally supported by NIH NIGMS P30GM110758.

## acronyms

MIC: minimum inhibitory concentration
BLI: biolayer interferometry
MST: microscale thermophoresis
SV AUC: sedimentation velocity analytical ultracentrifugation

## Notes

### Competing Interest Statement

The authors have declared no competing interest.

