## Supplemental Information - Fig. S1-S6, Table S1,S3,S4,S5 for "Phosphoarginine modulates oligomerization and repressor activity of mycobacterial ClpC2"

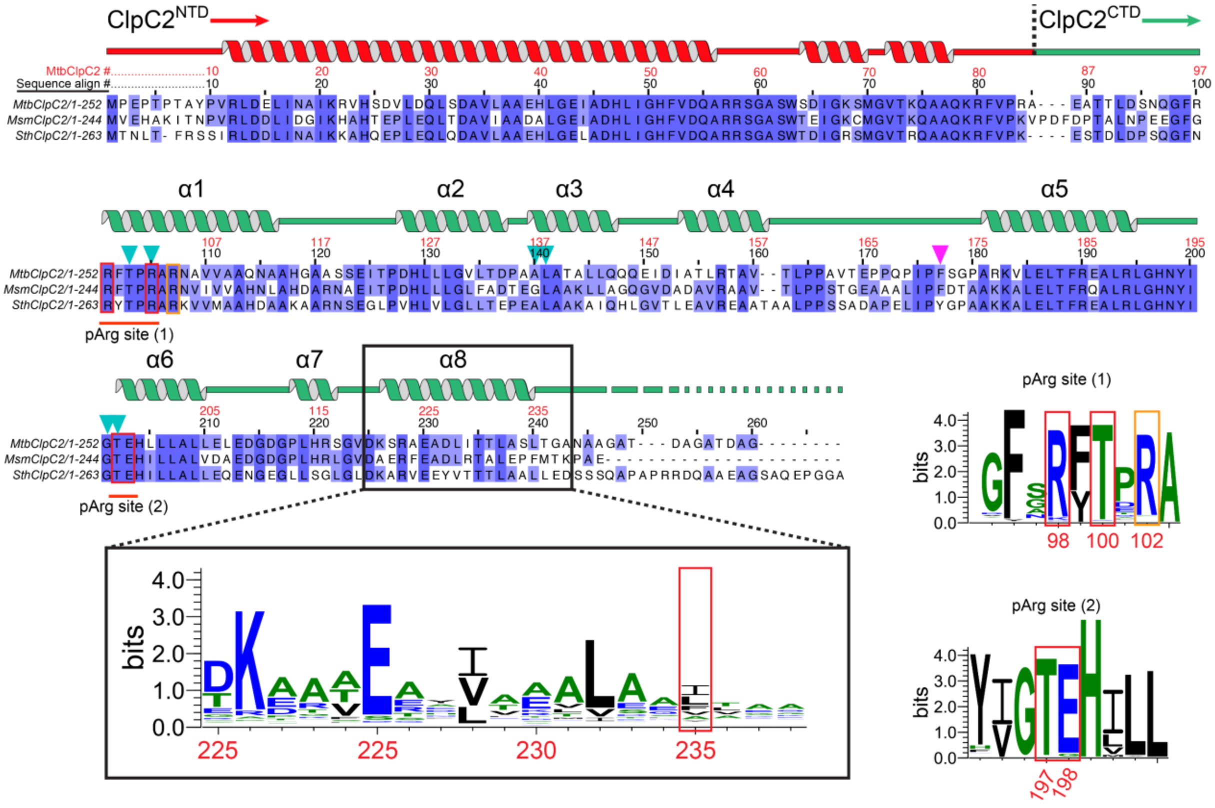

**Figure S1. Sequence conservation among ClpC2 orthologs.** A sequence alignment of ClpC2 orthologs from *M. tuberculosis*, *M. smegmatis*, and *S. thermoviolaceus* is shown, along with annotations of the domain organization and secondary structure. Sequence logos show conservation of the C-terminal α8 helix and the segments of the pArg binding site. Cyan triangles mark conserved residues that line the pocket where pArg / α8 bind. The magenta triangle marks a residue substitution that likely weakens Cym and Ruf binding in *S. thermoviolaceus.*

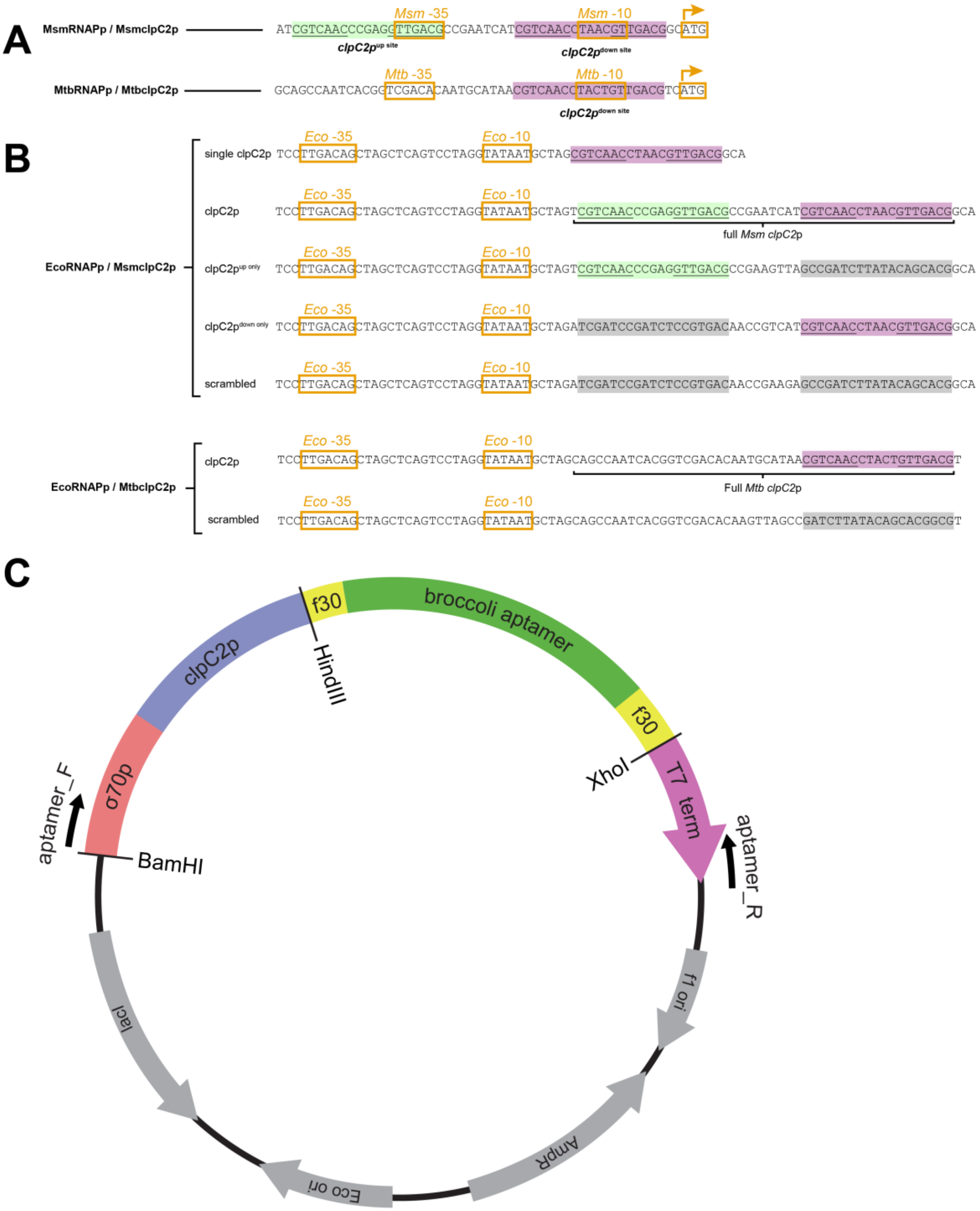

**Figure S2. Constructs incorporating ClpC2 operator sequences.** A) Genomic *clpC2* promoter regions from *M. smegmatis* and *M. tuberculosis*. B) Promoter and operator sequences used for *in vitro* translation assays, incorporating an *E. coli* σ70 promoter and downstream operator components. C) Architecture of the plasmid incorporating the transcription template construct.

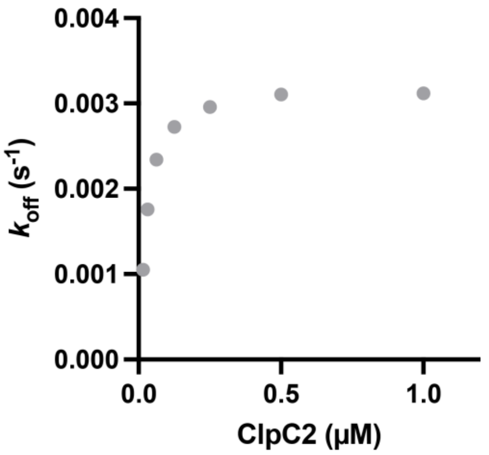

**Figure S3. Dissociation rates of *M. smegmatis* ClpC2 bound to *clpC2*o.** Varying concentrations of $^{Msm}$ClpC2 were allowed to bind to immobilized *clpC2o* (as in Fig. 3B). Biosensor tips were transferred to buffer, and dissociation trajectories were monitored. Observed dissociation rates are plotted against the initial $^{Msm}$ClpC2 concentration.

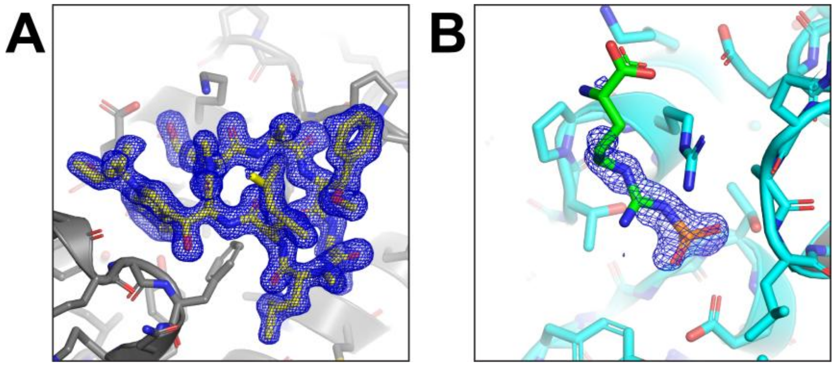

**Figure S4. Electron density maps covering ClpC2 ligands.** Electron density omit maps were calculated in the absence of A) CymA or B) pArg are shown at the respective ligand positions, contoured at 1.0 σ.

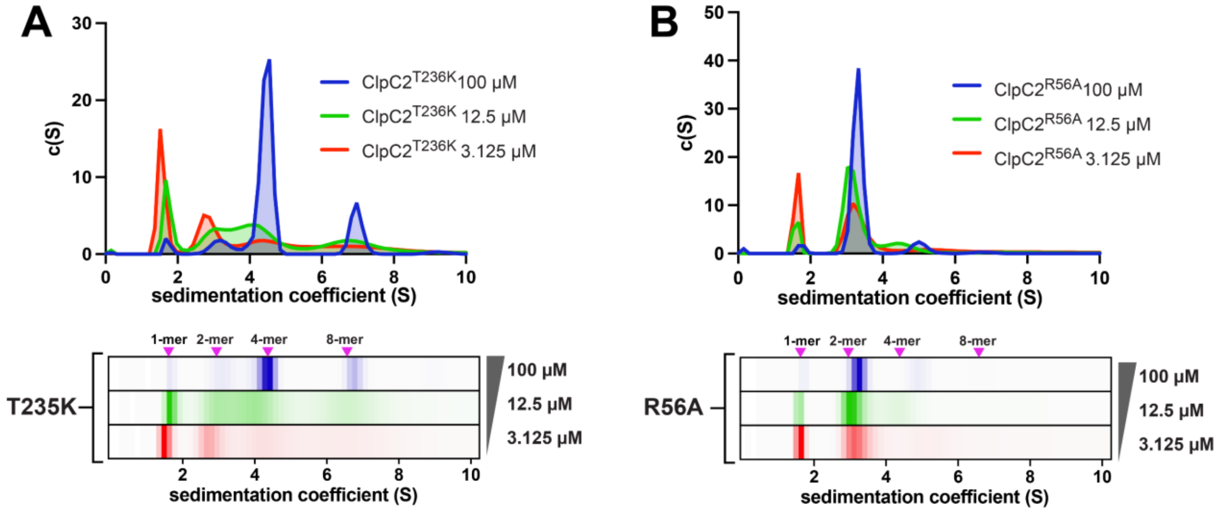

**Figure S5. Analytical ultracentrifugation of ClpC2 variants.** The indicated concentrations of MtbClpC2 variants were analyzed by velocity analytical ultracentrifugation and fit to sedimentation coefficient distributions shown as plots (upper) and one-dimensional heatmaps (lower) where darker colors correspond to higher *c(S)*. Mutations shift oligomer distribution towards lower-order species, compared to wild-type distributions in Fig. 2C. Arrows indicate expected sedimentation coefficients of the indicated species.

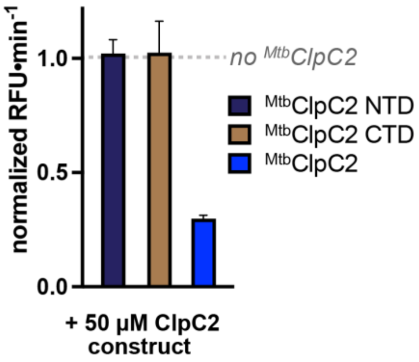

**Figure S6. ClpC2 NTD and CTD alone do not repress transcription.** *In vitro* assays monitored transcription of a F30-broccoli RNA aptamer from a template incorporating Mtb*clpC2o*. Transcriptional rate in the presence of MtbClpC2 (blue), MtbClpC2 N-terminal domain (NTD; black), or MtbClpC2 C-terminal domain (CTD; brown) is compared to the rate in the absence of MtbClpC2 (gray dashed line).

**Table S1. Binding affinities of compounds to ClpC2**

| | $^{Mtb}$ClpC2 | $^{Sth}$ClpC2 |
|---|---|---|
| | $K_D$ (nM) | $K_D$ (nM) |
| CymA (BLI) | 1.5 ± 0.1 | 86 ± 6 |
| Ruf (BLI) | 3.3 ± 0.3 | 110 ± 8 |
| Ecu (BLI) | n.b. | n.b. |
| pArg (MST) | 4700 ± 500 | 12000 ± 200 |
| pSer (MST) | n.b. | n.b |
| pThr (MST) | n.b. | n.b. |

Binding constants are reported ± standard fit error.
n.b.: no binding

**Table S2. Mass spectrometry data**. *see Supplemental Excel File*

**Table S3. Binding affinities of ClpC2 to immobilized DNA constructs**

| | $^{Mtb}$ClpC2 |
|---|---|
| | $K_D$ (nM) |
| Mtb $^{b}$*clpC2o* | 270 ± 20 |
| Mtb $^{b}$*clpc2o* scrambled | n.b. |

| | $^{Msm}$ClpC2 |
|---|---|
| | $K_D$ (nM) |
| Msm *pclpC2* | 10 ± 0.4 |
| Msm $^{b}$*clpC2o* up site | 240 ± 20 |
| Msm $^{b}$*clpC2o* down site | 250 ± 50 |
| Msm $^{b}$*clpC2o* scrambled | n.b. |

Binding constants are reported ± standard fit error.
n.b.: no binding

**Table S4. X-ray crystal structure collection and refinement statistics**

| PDB ID | **9P2P** | **9P39** | **9P4H** |
|---|---|---|---|
| Protein | $^{Sth}$ClpC2 CTD | $^{Sth}$ClpC2 CTD | $^{Sth}$ClpC2 CTD |
| Ligand | - | phosphoarginine | cyclomarin A |
| | **data collection statistics** | | |
| Space group | C121 | $P6_1$ | C121 |
| Cell dimensions | | | |
| x, y, z (Å) | 47.85, 46.06, 71.28 | 74.08, 74.08, 50.69 | 50.51, 44.06, 70.35 |
| $\alpha$, $\beta$, $\gamma$ (°) | 90, 105.53, 90 | 90, 90, 120 | 90, 109.36, 90 |
| X-ray source | CHESS 7B2 | CHESS 7B2 | NSLSII 17-ID-2 |
| Wavelength (Å) | 0.9686 | 0.9686 | 0.97934 |
| Max resolution (Å) | 1.41 | 1.30 | 1.55 |
| % Completeness | 95.8 | 92.0 | 96.6 |
| $CC_{1/2}$ | 0.999 | 1 | 0.998 |
| $\langle I/\sigma \rangle$ | 2.53 | 1.97 | 3.42 |
| | **refinement statistics** | | |
| Resolution range (Å) | 34.34 - 1.41 | 34.34 - 1.3 | 33.19 - 1.55 |
| R factor ($R_{free}$) | 0.189 (0.214) | 0.168 (0.19) | 0.148 (0.174) |
| Total/non H/solv. atoms | 2243 / 1110 / 140 | 2353 / 1169 / 118 | 2375 / 1192 / 164 |
| Ramachandran | | | |
| favored | 147 | 154 | 155 |
| allowed | 0 | 1 | 2 |
| outliers | 0 | 0 | 0 |
| Clashscore | 1 | 0 | 2 |
| RMSD bond length (Å) | 0.1 | 0.42 | 0.12 |
| RMSD bond angles (°) | 0.26 | 0.56 | 0.27 |

**Table S5. Primers used in this study**

| Name | Sequence | Modification | Purpose | |
|---|---|---|---|---|
| Aptamer_F | GATCGAGATCTCGATCCCGCGAAAT | - | Linearize IVT construct | IVT |
| Aptamer_R | GCAAAAAACCCCTCAAGACCCGTTTAGA | - | Linearize IVT construct | |
| Sig70_F | GATCCTTGACAGCTAGCTCAGTCCTAGGTATAA | - | Anneal for restriction cloning into IVT vector | |
| Sig70_R | CTAGCATTATACCTAGGACTGAGCTAGCTGTCAAG | - | Anneal for restriction cloning into IVT vector | |
| MsmclpC2p_F | TGCTAGTCGTCAACCCGAGGTTGACGCCGAATCATCGTCAACCTAACGTTGACGGCA | - | Anneal for restriction cloning into IVT vector | |
| MsmclpC2p_R | AGCTTGCCGTCAACGTTAGGTTGACGATGATTCGGCGTCAACCTCGGGTTGACGA | - | Anneal for restriction cloning into IVT vector | |
| MsmclpC2p_downonly_F | TGCTAGATCGATCCGATCTCCGTGACAACCGTCATCGTCAACCTAACGTTGACGGCA | - | Anneal for restriction cloning into IVT vector | |
| MsmclpC2p_downonly_R | aGCTTGCCGTCAACGTTAGGTTGACGATGACGGTTGTCACGGAGATCGGATCGAT | - | Anneal for restriction cloning into IVT vector | |
| MsmclpC2p_uponly_F | TGCTAGTCGTCAACCCGAGGTTGACGCCGAAGTAGCCGATcTTAtACaGCacGgcGA | - | Anneal for restriction cloning into IVT vector | |
| MsmclpC2p_uponly_R | aGCTTCgcCgtGCtGTaTAAgATCGGCTACTTCGGCGTCAACCTCGGGTTGACGA | - | Anneal for restriction cloning into IVT vector | |
| MsmScrambled_both_F | TGCTAGATCGATCCGATCTCCGTGACAACCGGTAGCCGATcTTAtACaGCacGgcGA | - | Anneal for restriction cloning into IVT vector | |
| MsmScrambled_both_R | aGCTTCgcCgtGCtGTaTAAgATCGGCTACCGGTTGTCACGGAGATCGGATCGAT | - | Anneal for restriction cloning into IVT vector | |
| FullMtbC2P_fragF | TGCTAGcagccaatcacggtcgacacaatgcataacgtcaacctactgttgacgt | - | Anneal for restriction cloning into IVT vector | |
| FullMtbC2P_fragR | acgtcaacagtaggttgacgttatgcattgtgtcgaccgtgattggctgTCGa | - | Anneal for restriction cloning into IVT vector | |
| Up_downscrMtbC2P_fragF | TGCTAGcagccaatcacggtcgacacaaGTAGCCGATcTTAtACaGCacGgcGt | - | Anneal for restriction cloning into IVT vector | |
| Up_downscrMtbC2P_fragR | aCgcCgtGCtGTaTAAgATCGGCTACttgtgtcgaccgtgattggctgTCGa | - | Anneal for restriction cloning into IVT vector | |
| **Name** | **Sequence** | **Modification** | **Purpose** | Octet |
| bio-MsmclpC2p_F | TCGTCAACCCGAGGTTGACGCCGAATCATCGTCAACCTAACGTTGACGGC | 5' biotin-GG | Anealing for octet with 5'biotin tag | |
| MsmclpC2p_R | GCCGTCAACGTTAGGTTGACGATGATTCGGCGTCAACCTCGGGTTGACGA | - | Anealing for octet with 5'biotin tag | |
| bio-MsmclpC2p_downOnly_F | ATCGATCCGATCTCCGTGACAACCGTCATCGTCAACCTAACGTTGACGGC | 5' biotin-GG | Anealing for octet with 5'biotin tag | |
| MsmclpC2p_downOnly_R | GCCGTCAACGTTAGGTTGACGATGACGGTTGTCACGGAGATCGGATCGAT | - | Anealing for octet with 5'biotin tag | |
| bio-MsmclpC2p_upOnly_F | TCGTCAACCCGAGGTTGACGCCGAAGTAGCCGATcTTAtACaGCacGgcG | 5' biotin-GG | Anealing for octet with 5'biotin tag | |
| MsmclpC2p_upOnly_F | CgcCgtGCtGTaTAAgATCGGCTACTTCGGCGTCAACCTCGGGTTGACGA | - | Anealing for octet with 5'biotin tag | |

| | | | |
|---|---|---|---|
| bIVT_amp_F | GATCGAGATCTCGATCCCGCG | 5' biotin-GG | Amplifying MtbclpC2o sequences |
| bIVT_amp_R | cgtctcccacatacacatggcaa | - | Amplifying MtbclpC2o sequences |
| bio-Scrambled_F | ATCGATCCGATCTCCGTGACAACCGGTAGCCGAT cTTAtACaGCacGgcG | 5' biotin-GG | Anealing for octet with 5'biotin tag |
| Scrambled_R | CgcCgtGCtGTaTAAgATCGGCTACCGGTTGTCA CGGAGATCGGATCGATy | - | Anealing for octet with 5'biotin tag |
